# Enhanced chemotaxis through spatially-regulated absolute concentration robustness

**DOI:** 10.1101/2021.07.10.451673

**Authors:** Debojyoti Biswas, Sayak Bhattacharya, Pablo A. Iglesias

## Abstract

Chemotaxis, the directional motility of cells in response to spatial gradients of chemical cues, is a fundamental process behind a wide range of biological events, including the innate immune response and cancer metastasis. Recent advances in cell biology have shown that the protrusions that enable amoeboid cells to move are driven by the stochastic threshold crossings of an underlying excitable system. As a cell encounters a chemoattractant gradient, the size of this threshold is regulated spatially so that the crossings are biased towards the front of the cell. For efficient directional migration, cells must limit undesirable lateral and rear-directed protrusions. The inclusion of a control mechanism to suppress these unwanted firings would enhance chemotactic efficiency. It is known that absolute concentration robustness (ACR) exerts tight control over the mean and variance of species concentration. Here, we demonstrate how the coupling of the ACR mechanism to the cellular signaling machinery reduces the likelihood of threshold crossings in the excitable system. Moreover, we show that using the cell’s innate gradient sensing apparatus to direct the action of ACR to the rear, suppresses the lateral movement of the cells and that this results in improved chemotactic performance.

## 1 Introduction

For a chemical reaction to ensue, two physical processes must take place. First, the interacting molecules must come together. The rate at which this process takes place is dictated by the diffusivity of the molecules which, at the microscopic level, is governed by the Brownian motion of the particles. Once the molecules are within a given interaction radius, a chemical conformational change takes place. This process is also stochastic, as it is usually viewed as overcoming a stochastic potential well [1, 2]. The effect of this randomness on the concentration of the biochemical species is quite acute when the copy number of the interacting molecules is small, such as genetic regulatory networks. Such systems have attracted considerable theoretical and experimental attention [3–5]. However, stochastic effects can also influence cellular behavior even when the number of molecules is large. The migration of amoeboid cells is one such example.

The social amoeba *Dictyostelium discoideum* lives in the soil where it forages for nutrients, typically bacteria [6, 7]. To find food, it is continuously in motion which takes the form of actin-filled protrusions known as pseudopods (literally “false feet”). In migrating cells, these protrusions appear randomly around the cell perimeter but have reproducible characteristics, such as their size, frequency and lifetime [8, 9]. These cells also can direct pseudopods in the direction of their prey, by using chemical traces emanating from their target, a process known as chemotaxis. Specifically, bacteria secrete folic acid which binds to cell-surface receptors that are specific to this chemical. Through a complex but sophisticated method, the amoebae can discern the direction of the gradient, and increase the local probability of extending a pseudopod in that direction, while simultaneously lowering the probability of pseudopods away from the source. This spatially biases the extension of pseudopods, resulting in the directed motion that enables the cell to survive.

The characteristics of the pseudopods for cells that are migrating randomly or following a chemoattractant gradient are quite similar [8], suggesting that there is a common mechanism regulating the formation of a pseudopod, and that chemoattractant receptor occupancy is used solely to spatially guide this process. During the past fifteen years, it has been increasingly clear that the appearance of these extensions in *Dictyostelium* cells is regulated by an excitable system [10, 11]. The movement of cells using an excitable system is not limited to these amoebae. The existence of an excitable cortex is highly conserved and has been detected in nematodes [12], *Xenopus* frog eggs [13] as well as mammalian white blood cells (neutrophils [14–16], mast cells [17] and macrophages [18]), cultured neurons [19] and cancer cells [20, 21].

An excitable system is a class of nonlinear dynamical systems that has several features [22]. Having a single stable equilibrium, small perturbations about this equilibrium elicit small-scale responses. However, whenever the perturbations are sufficiently large, they produce a large-scale deviation away from the equilibrium before the state returns to its prestimulus level. While the system is undergoing this trajectory, it exhibits a refractory period during which no further activation is possible. Both the size and length of this excursion are also characteristic of the system, which can be thought to possess a threshold for activation [23]. Stochasticity can induce the system to cross the threshold triggering a large-scale excursion. The rate at which this happens is governed by the ratio of the size (standard deviation) of the noise relative to that of the threshold [24, 25].

It has been hypothesized that the extension of pseudopods represents the crossing of this firing threshold [26–28] In a randomly migrating cell, the size of the threshold is uniform throughout the cell, and hence the probability that the threshold is crossed leading to the extension of a pseudopod is spatially, uniformly distributed around the perimeter. However, in the presence of a gradient of receptor occupancy, the threshold is spatially modulated. At the site closest to the chemoattractant source, the threshold is lowered, making it easier to cross and hence increasing the likelihood that pseudopods are extended. Away from the source, the threshold is increased reducing the likelihood that pseudopods will form. The resulting biased, excitable network can move cells in the direction of the chemoattractant [27, 29–33]. To date, there is ample evidence of this model of cell movement, from the existence of sub/supra-threshold behavior and refractory periods [34,35], to the resultant traveling waves that are characteristic of excitable media [14, 26, 36–39]. Recently, it has been possible to alter this threshold synthetically, leading to large changes in the behavior of the dynamical system [21, 40–43].

In the presence of shallow gradients, the raising of the threshold at the rear of the cell may not be sufficient to abrogate completely the extension of pseudopods. These rear-directed pseudopods would have the net effect of slowing overall migration towards the chemoattractant source. There are two potential ways to limit these undesirable protrusions. In theory, one could engineer a mechanism to increase the threshold at the back further. Alternatively, one could design a control system that provides some control over the variance in the fluctuations of specific molecules that trigger cell movement. In this paper, we consider the latter through a mechanism known as absolute concentration robustness (ACR) [44–46]. ACR comes about through the binding of a target species in a specific reaction scheme, which ensures that the steady-state concentration of the target remains robustly stable to fluctuations in its environment. In this study, we show how ACR can be used to achieve control over the levels of critical lipids in the cellular signaling network which in turn directly affects the firing rate of the underlying excitable system. Together, this creates a control scheme for faster and more efficient directed migration.

The rest of the paper is organized as follows. In Section 2, we consider some mathematical preliminaries. We describe a simplified mathematical model of an excitable reaction network that closely recreates the dynamics of the system regulating actin protrusions. We also present equations describing the mathematical model of the ACR regulator. In Section 3, we demonstrate how ACR can be used to suppress excitable network activity and describe the signaling system used to sense and interpret chemoattractant gradients. Following that, we present our main findings: how the inclusion of the ACR system can reduce the variance of the stochastic perturbations. Further, we couple this ACR system with a spatially heterogeneous network to focus its activity at the rear of migrating cells. By incorporating a simple model of cell motion, we demonstrate, through simulation, enhanced chemotaxis towards the chemoattractant source. Finally, in Section 4, we present some conclusions.

## 2 Preliminaries

### 2.1 Mathematical model of the excitable network regulating motility

The seminal work of FitzHugh [47] and Nagumo [48] showed that it is possible to describe the dynamics of excitability through a two-state activator-inhibitor network. The activator (*u*) incorporates a positive feedback loop that allows it to stimulate its production. The inhibitor (*v*), whose dynamics occur on a slower time scale, provides negative feedback to the activator. In the classical description of cellular excitability, the variables represent membrane potentials, which may be negative. In our case, excitability comes about because of activities and concentrations of various interacting molecules, which are non-negative variables. To ensure that this is the case, we considered a two-dimensional activator-inhibitor model given by [23] :

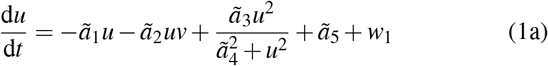

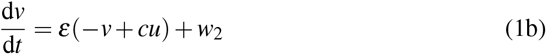

This model captures the dynamics of excitable systems while ensuring that all species concentrations remain non-negative. The essential nonlinearity in the activator dynamics in Eqn. 1a is incorporated through a co-operativity term with Hill coefficient two. The inhibitor dynamics operate at a slower time scale owing to the parameter *ɛ* ≪ 1 in Eqn. 1b. The terms *w*_1_ and *w*_2_ represent the contributions of any external signals to the system.

In the context of cell migration, recent evidence has suggested that the autocatalytic behavior of the activator is likely achieved via a third species through mutual inhibition [49]. This is based on the spatial segregation of biochemical species seen during chemotaxis, where some species accumulate towards the source (front), while others accumulate at the opposite end of the cell (back) [50]. This mutually exclusive segregation is characteristic of mutual inhibition [51]. To accommodate these new findings, we modified the activator-inhibitor system as a three-dimensional system, renaming the variables *F*, *B*, and *R* for “front”, “back” and “refractory,” respectively (Fig. 1A).

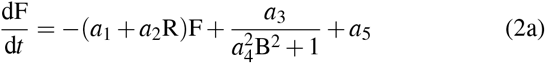

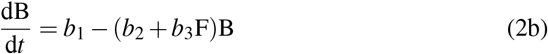

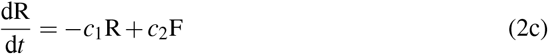

**Figure 1:**
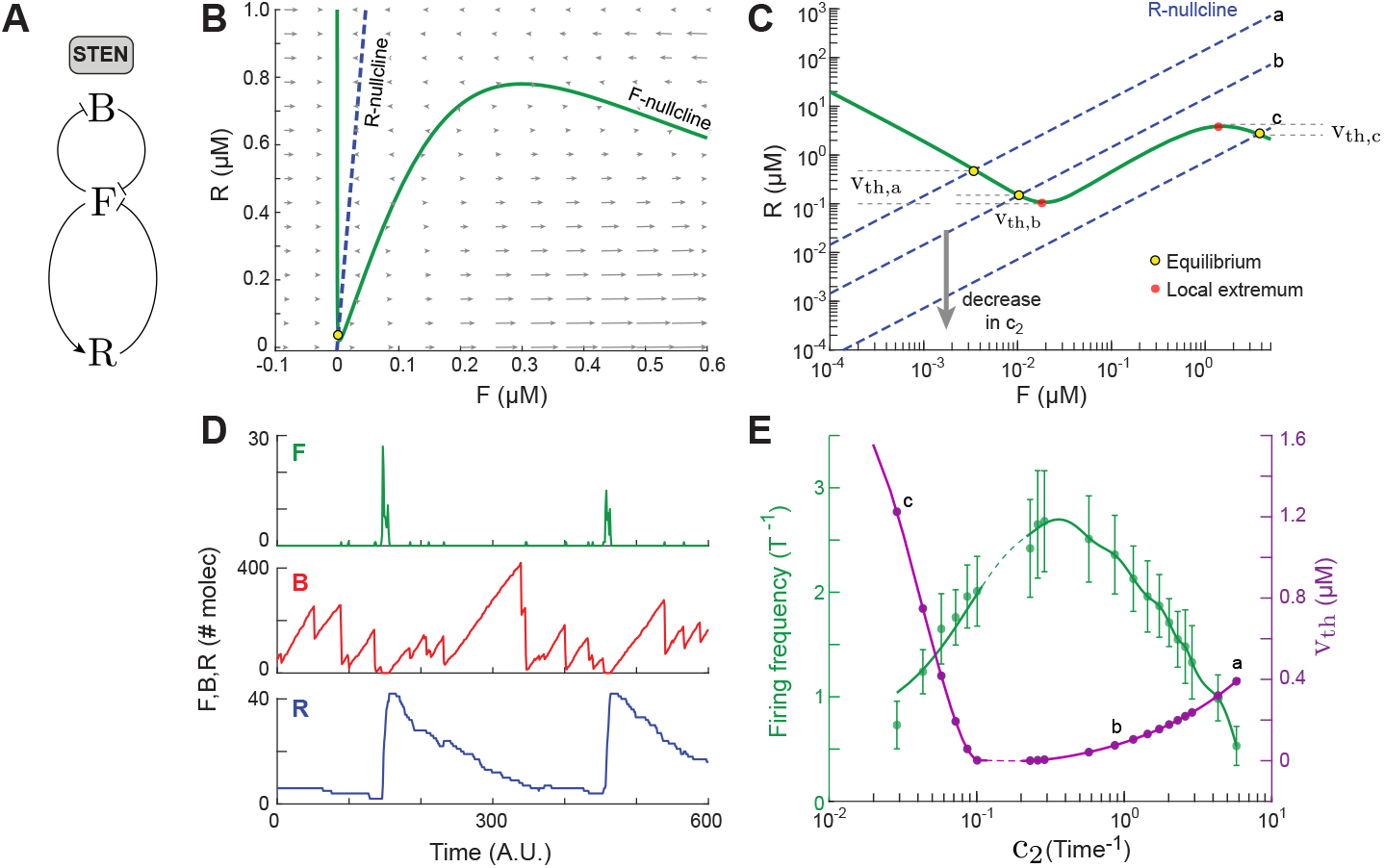
Excitable dynamics. (A) Schematic of the *signal transduction excitable network* (STEN) showing the entities F, B and R and their interactions. (B) F-R phase portrait showing the respective nullclines (F: green solid, R: blue dashed) and the equilibrium point (yellow circle). The arrows represent the magnitude and direction of velocities (Ḟ, Ṙ) in the field. (C) Lowering of F-dependent activation rate of R (*c*_2_) results in a change in position of the R-nullcline (*a* → *b* → *c*) as well as in *v*_th_. The local extrema of the F-nullcline are denoted as red circles. (D) Typical time profiles of F, B and R. Parameter values used in simulation: *a*_1_ = 2, *a*_2_ = 20, *a*_3_ = 240, *a*_4_ = 120, *a*_5_ = 0.04, *b*_1_ = 3, *b*_2_ = 0.006, *b*_3_ = 240, *c*_1_ = 0.08 and *c*_2_ = 5.76. (E) Plots of the firing frequency and threshold (*v*_th_) as a function of parameter *c*_2_. The firing frequency was measured by counting the number of firings in a time window of 600 A.U. The dashed section corresponds to the parameter space where *v*_th_ is undefined following the definition from Eqn. 4.

We refer to this system of equations as the *Signal Transduction Excitable Network* (STEN) [10].

Under the quasi-steady-state assumption on Eqn. 2b as the second reaction approaches steady-state: B = *b*_1_*/*(*b*_2_ + *b*_3_F), and the remaining two equations assume the form of Eqn. 1. The dynamics of this system can be better visualized using phase space (Fig. 1B). Whereas the F-nullcline is cubic and displays an inverted “N-shape,” the R-nullcline is linear. Changes in the parameter *c*_2_, in Eqn. 2c, alter the slope of the R-nullcline (which thus gets translated in log scale) as shown in a zoomed-out view of the phase space (Fig. 1C). These variations could affect system behavior as they may alter the stability of the existing equilibrium, or result in the emergence of multiple equilibria [23]. The normal mode of operation in chemotaxis is at position “a”, where a stable equilibrium exists to the left of the minimum of the cubic nullcline. If an external input or intrinsic noise in the system is sufficient to displace the state beyond this minimum, then the state (F, R) undergoes a large excursion (firing) in phase space, creating a spike in time observed in both the F and R states (Fig. 1D). This is followed by a refractory period, during which no further firings are possible, as the inhibitor (R) decays back to equilibrium. B shows a complementary profile to F [49] and thus reaches a minimum whenever a firing takes place.

We now briefly demonstrate what parameters control the firings of the excitable network. According to Kramer’s theory [1, 2], the relationship between firing frequency, *f*, and the noise level is given by

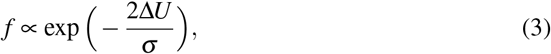

where Δ*U* is the height of a potential barrier and *σ* is the noise variance. This potential barrier is related to the activation threshold of the excitable system, *v_th_*, that can be described as (Fig. 1C):

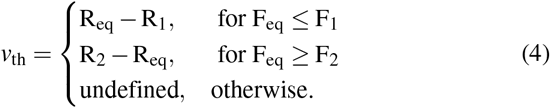

Here, (F_1_, R_1_) and (F_2_, R_2_) are the local minimum and maximum, respectively of the cubic nullcline (red markers in Fig. 1C). In this definition of *v_th_*, we only consider parameter values that give rise to a single, stable equilibrium and have excluded cases of unstable and/or multiple equilibria (F_eq_ ∈ (F_1_, F_2_)). Note, that the parameter space that gives rise to multiple equilibria in our system is small.

Decreasing *c*_2_ (e.g., to point “b” in Fig. 1C) lowers the threshold level which in turn increases the firing rate until (F_eq_, R_eq_) reaches (F_1_, R_1_) (Fig. 1E). Beyond F_1_, a bifurcation occurs and the equilibrium is rendered unstable resulting in a limit cycle. Upon decreasing *c*_2_ further, another stable equilibrium is created at (F_2_, R_2_). As *c*_2_ is lowered even further, the threshold *v*_th_ increases resulting in lower firing rates. Thus, the firing frequency shows a biphasic relationship with the activation threshold of the network (Fig. 1E).

### 2.2 Protrusions in spatial simulations

So far, we have used temporal simulations of the excitable network to explain how noise-induced firings occur. These firings are posited to lead to cellular protrusions and hence cell motility. To describe the effect on *directional* motility, however, requires that we consider a spatial dimension. To this end, we extend our excitable system in one spatial dimension which represents the perimeter of a circular cell. In this spatial model, we allow the reacting species to diffuse isotropically. Plotting the activity as a function of time and space (*x* and *y*-axes, respectively) gives rise to the kymographs of Fig. 2. In these kymographs, the firing of the system is seen as an initial trigger that spreads in space (along the cell perimeter) in the form of a traveling wave (bright F and R; dark B) [52]. The F and B kymographs show complementary patterns whereas the R kymograph is similar to that of F, but more diffused in time owing to the slower time scale in the refractory equation.

**Figure 2:**
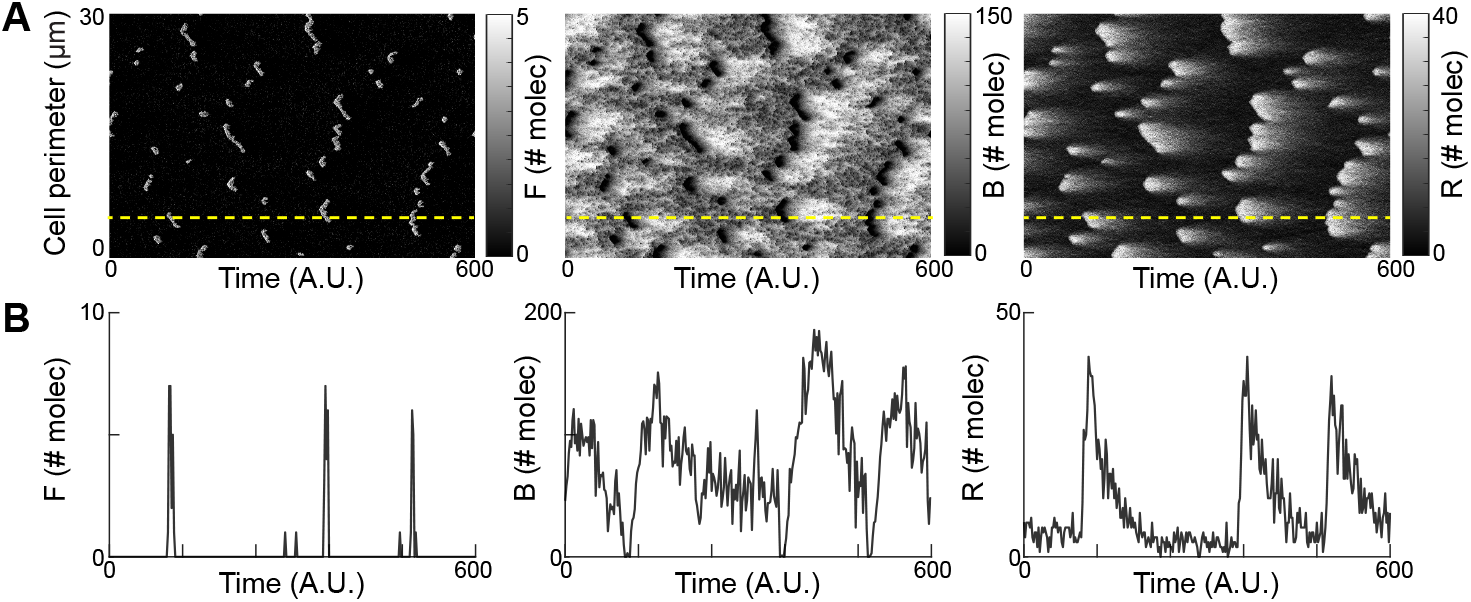
Spatiotemporal simulation of excitable dynamics. (A) Kymographs of F, B and R. Brighter regions represent higher molecule numbers. The F and B kymographs show complementary patterns whereas the R kymograph is similar to that of F but more diffused in time. (B) Temporal profiles of F, B and R at the location marked by the yellow dashed lines in panel A.

In cells, these spatiotemporal waves correspond to protrusions on the cell membrane. A line scan through the kymographs shows the activity in Fig. 2B as a function of time at one point on the cell perimeter (yellow dashed line in Fig. 2A). These profiles match the previous plots in Fig. 1D. These kymographs closely resemble similar plots of cell protrusions obtained in experiments [21, 40, 49].

### 2.3 Absolute concentration robustness

Per Eqn. 3, the rate at which activity is triggered in an excitable system depends on both the size of the threshold as well as the size of the noise. Having seen how the threshold affected the firings, we now consider the role of the noise and, in particular, how absolute concentration robustness (ACR) could be used to limit these firings. To illustrate the essential features of ACR, we consider the following toy model [44]. This model consists of two biochemical species: the target species, M, whose concentration is to be regulated, and N, which achieves this regulation. The two species obey the following mass action reaction scheme (Fig. 3A, top)

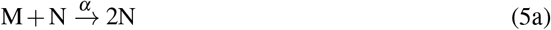

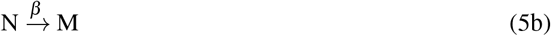

**Figure 3:**
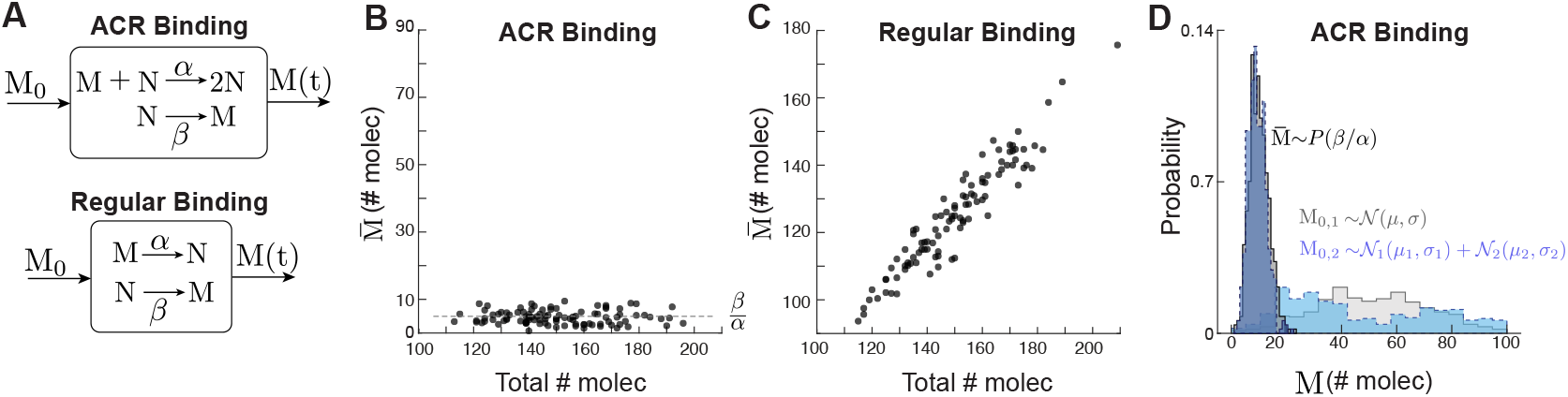
Comparison between ACR and regular binding. (A) Schematic of ACR and regular binding with species M, N and their respective rate constants. M_0_ and M(t) are the initial condition and the system output, respectively. (B, C) Steady-state concentrations of 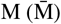 following stochastic simulations (using SSA) of ACR (B) and regular binding (C) for varying total number of molecules. (D) Steady-state distributions of M following stochastic simulations of ACR for two different initial distributions, 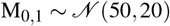 and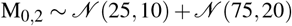. Simulations used *α* = 0.1 and *β* = 0.5.

We assume that the total number of molecules is conserved (i.e. M + N = constant = T). Solving for the steady-state concentrations of the species leads to two possible equilibria:

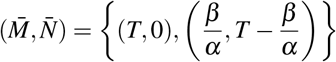

where the first is unstable and the second stable. Thus, as long as the initial steady-state avoids the unstable equilibrium, the steady-state concentration of M is constant. This depends only on the reaction rates, *α* and *β* (Fig. 3B), and not on the concentration of the binding species. This is in contrast to the case of regular binding reactions (Fig. 3A, bottom):

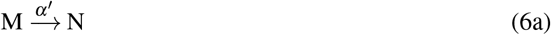

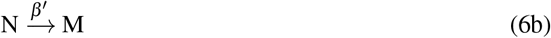

where the steady-state is given by

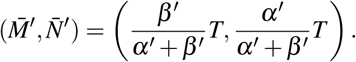

Unlike ACR binding, here the steady-state of M is a linear function of the total concentration (Fig. 3C).

It follows that, in a deterministic setting, the ACR scheme provides a robust mechanism for maintaining a constant concentration of a particular reactant. Our main interest, however, is in controlling concentration variations. As shown by Anderson et al. [45], the ACR scheme enables the system to control the variance of the target species. In particular, for a large total number of molecules, the stationary distribution of M assumes a Poisson distribution with parameter *β /α.* The advantage of a Poisson process is that variance is now equal to the mean:

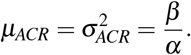

The same is not necessarily true for normally distributed systems where the mean and variance are independent parameters. For large means, Poisson fluctuations are small relative to the mean. Thus, in the case of a Poisson process, if we have a method to control the mean of the system, we can automatically put a restriction on the variance. Fig. 3D illustrates the results of a stochastic simulation of the ACR scheme assuming two different initial distributions, a unimodal Gaussian distribution given by 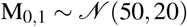 (gray) and a bimodal distribution given by the sum of two Gaussians: 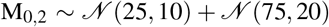 (blue). In both cases, the steady-state distribution converges to a Poisson distribution with mean and variance ≈ *β/α* = 5. In this way, the ACR scheme grants robust control over both the mean and fluctuation level of the process.

### 2.4 Stochastic simulations

Although, in this paper, we have described all systems using reactions and their corresponding differential equations, all our simulations are, in fact, based on a stochastic description of the systems. This requires that reactions be described by propensity functions. Descriptions of the propensity functions for all simulations and the corresponding parameters are listed in Table 1.

**Table 1:**
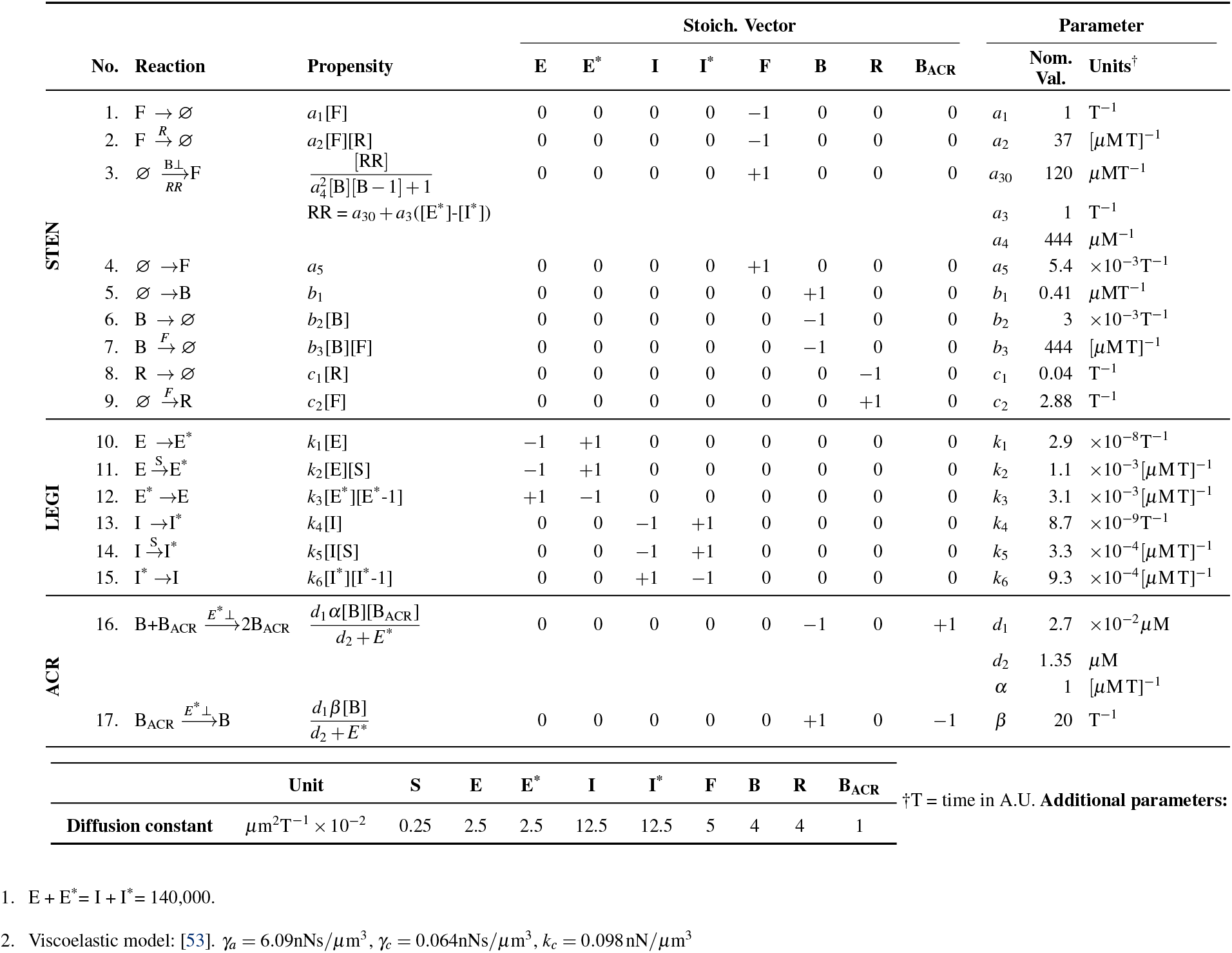
Parameters used in the stochastic simulations.

For these stochastic simulations, we adopted two methods. First, for homogeneous systems, we used Gillespie’s stochastic simulation algorithm (SSA) [54]. Briefly, in the SSA, at each time point in the simulation, two random numbers are used to find the identity of the next reaction and the time at which it is to takes place. In the limit, the distribution obtained from these SSA simulations exactly recreates the probability distribution of the chemical master equation that describes the system. Second, for spatially heterogeneous systems, we used the reaction-diffusion master equation (RDME). Here, the spatial volume is discretized into smaller voxels. Within each subvolume, the reactions are assumed to follow the statistics of a well-stirred system. Additionally, diffusion is modeled as a first-order reaction transferring molecules from one voxel to another one. To implement the RDME, we used the software package URDME [55], which allows for unstructured lattices and uses the Next Sub-volume Method [56] to simulate the system. For our spatiotemporal simulations, we assumed a one-dimensional cell perimeter of length 30 *μ*m with periodic boundary conditions (roughly equivalent to the perimeter of a circular cell of radius 5 *μ*m). The domain was subdivided into 601 equally-sized voxels.

## 3 Main Results

### 3.1 Controlling the firing frequency of the excitable system through ACR

As argued above, membrane protrusions are initiated by the firings of an excitable network, and the frequency of these events is controlled by the size of the noise relative to that of the threshold. The results of Section 2.3 suggest that this noise could be regulated through the inclusion of an ACR mechanism in the excitable network. For directed migration, we want to inhibit protrusions at the rear of the cell (i.e. away from the chemoattractant source). Thus, we are primarily interested in regions where the concentration of “back” (B) molecules is high. Thus, we couple the ACR scheme to the excitable network Eqn. 2 through state B. In particular, we posit the existence of a controlling variable B_ACR_ that binds to B according to the ACR scheme:

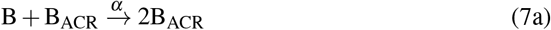

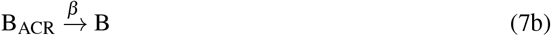

Thus, Eqn. 2b becomes:

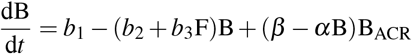

where B_ACR_ obeys:

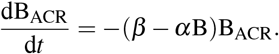

Note, that from the latter, in the steady-state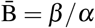 which forces the concentration of F to:

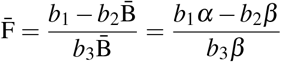

In particular, note that this is independent of the parameters in Eqn. 2a and Eqn. 2c.

To explore the effect of this scheme on the firings of the excitable system, we used spatial simulations, shown in Fig. 4. For the first half of the simulation, the system followed the dynamics of the regular F, B, R system. Thereafter, the ACR scheme was turned on by the incorporation of the B_ACR_ molecule. The kymographs of the F and B molecules show how the concentrations of both molecules decreased upon the incorporation of ACR, reaching a steady state after approximately 200 time steps (Fig. 4A). Once at steady-state, the ACR controller completely suppressed firings. Additionally, as shown in Fig. 4B, the variance of the B molecule was also suppressed.

**Figure 4:**
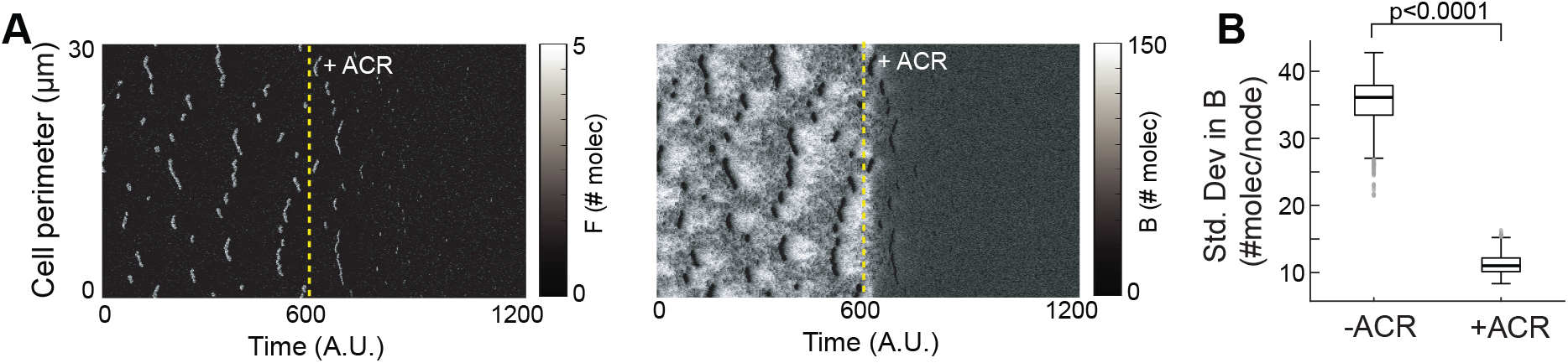
ACR controller suppresses firing. (A) Kymographs of F (left) and B (right) showing the effect of the ACR controller. ACR controller was manually turned on at t = 600 A.U. (dashed yellow line) and was able to suppress the activities within 200 time units. (B) Boxplot showing the change in the standard deviation of B computed over time and space (voxels) before and after the ACR controller was on. Simulations used *α* = 0.2 and *β* = 4.

### 3.2 Local-excitation, global inhibition based directed cell migration

The spatial simulations of the excitable network of Fig. 2 and Fig. 4 recreated patches of F and R molecules. Along with the downstream effectors, these patches lead to the formation of protrusions. However, these patches do not show any preferential direction. Thus, these kymographs represent the random, undirected movement of cells. To achieve directional sensing, it is important that an external gradient bias the location of these firings. In cells, this is achieved through a mechanism known colloquially as LEGI (local-excitation, global inhibition) [26].

The basic scheme is represented in Fig. 5A. In response to a chemoattractant signal, S, two species are activated that provide complementary excitation (E*) and delayed inhibition (I*) to a third species: the response regulator (RR). What allows this system to interpret chemical gradients is the difference between the diffusion properties of the different species [57]. Specifically, whereas the excitation molecule and the response regulator are mostly local, indicative of low diffusivity, the inhibition molecule is mostly global because of high diffusivity. In this study, we implement this system using the following reactions:

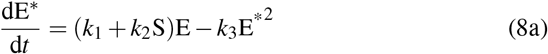

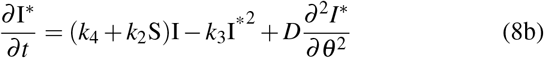

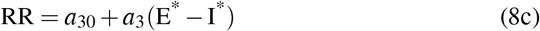

**Figure 5:**
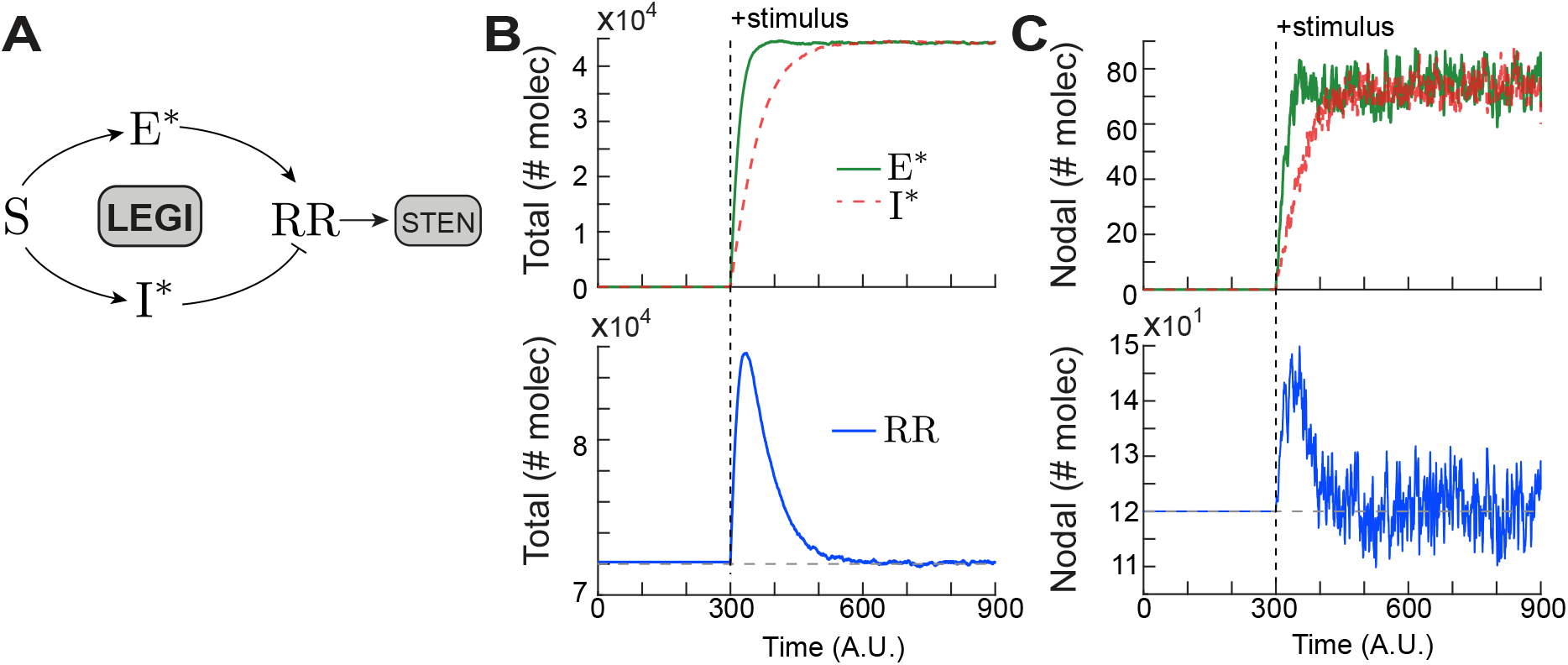
LEGI-STEN coupling achieves adaptation in response to global stimulus. (A) The schematic of LEGI shows the entities: S, E*, I*, RR and their respective interactions. The coupling between LEGI and STEN is coordinated through RR which represents the difference between E*and I*. (B,C) Total (B) and nodal (C) profiles of E*, I*and RR, before and after a global stimulus (dashed line).

The first terms in Eqn. 8a and Eqn. 8b denote the stimulus-independent activation of E and I. The quadratic term in Eqn. 8a represents degradation, mimicking the recombination G-protein heterotrimer components [58, 59]. Temporally, the system takes the form of an incoherent feedforward network. The dynamics of E*and I* assume that the inhibitor is delayed relative to the excitation molecule (Fig. 5B, top). Thus, when the exciter is activated in response to a stimulus, there is a brief period during which the inhibitor has yet to catch up, resulting in a rise in RR (Fig. 5B, bottom). As the mean levels of E and I converge at steady-state, the response regulator returns to its prestimulus level, ensuring adaptaion to a spatially homogeneous stimulus [15], as is characteristic of incoherent feedforward networks.

The LEGI mechanism can be used to stimulate an excitable network, for example, by making *a*_5_ in Eqn. 2a proportional to the level of RR. This causes increases in RR levels to lower the firing threshold of the excitable system. Thus, the transient pre-adaptation response of the LEGI response regulator, due to a stimulus, induces global firings of the excitable system. Eventually, as RR adapts, these firings subside and the system returns to its prestimulus state. This is the basis of the LEGI-biased excitable network that is believed to guide cell migration [10, 26, 28].

The plots of Fig. 5B show the behavior of E*, I* and RR summed over the whole spatial domain. In the spatial simulations, we focused on one randomly-selected voxel, to see more easily the effect of stochastic fluctuations in the constituents of LEGI (Fig. 5C). After the adaptation, the fluctuations in the E*and I*profiles give rise to a noisy response regulator (Fig. 5C). This noise input from RR contributes to the increased fluctuations in STEN - resulting in random firings.

Now, we focus on the direction-biasing capability of LEGI. This capablity arises from the difference in diffusion levels of the exciter and the inhibitor, as mentioned earlier. As shown in the kymographs and respective line scans of Fig. 6, we assume that the cell experiences a chemoattractant gradient resulting in a spatial bias in the concentration of stimulus S, creating front and back regions. Fig. 6B shows the effect of the gradient on the LEGI-STEN system. Over time, RR reaches a state where, in the region facing the gradient (line scan through “a”), it is above its basal level, and in regions away from the gradient (line scan through “b”), it is below (Fig. 6C). Because RR serves as the input to the excitable network, the elevated level of RR induces more activity in F in the front of the cell. Consequently, after the chemoattractant was presented, the patches in the region with the highest level of S (marked “a”) were enhanced, while those in the regions near it (marked “b”) saw a decline in overall activity. Far away (position marked “c”) there was no appreciable change, largely because the inhibitor had not diffused there. Eventually, as the LEGI system reached its steady-state, this region too saw diminished activity similar to that of point “b”. Thus, LEGI created a spatial activity bias in the cell, which would be reflected as the spatial segregation or self-organization of biochemical species on the cell membrane [50, 60].

**Figure 6:**
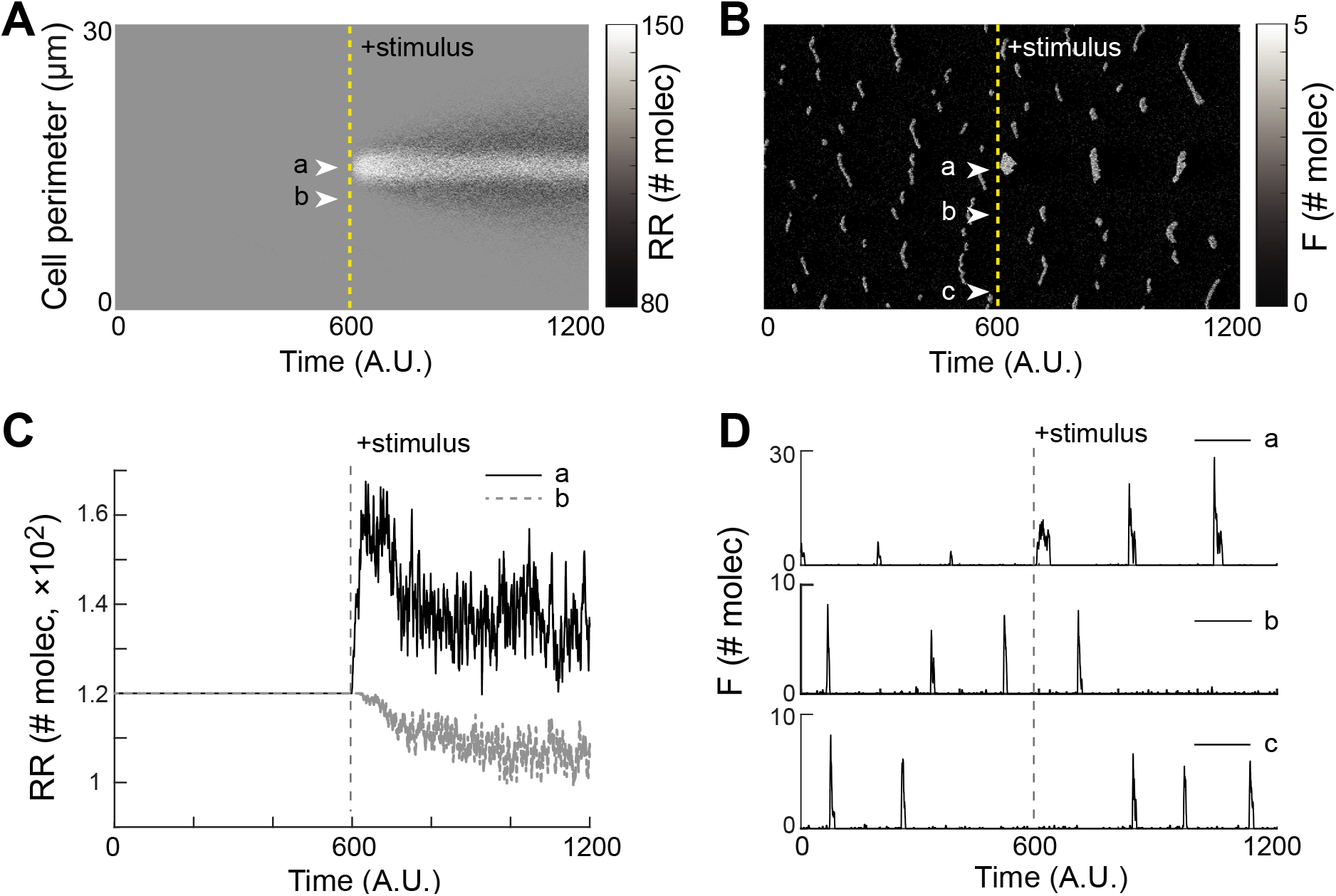
Gradient sensing by the coupled LEGI-STEN system. Kymographs of RR (A) and F (B) in response to a gradient stimulus applied at *t* = 600 A.U., along with the respective time profiles of RR (C) and F (D) from line scans at locations denoted by arrows, and marked a, b and c).

### 3.3 Enhanced chemotaxis: combining LEGI and ACR

The biased local-excitation, global inhibition input to the excitable system ensured increased activities towards the source of chemoattractant. However, occasional unwanted firings at the back of the cell are also observed which cause undesirable protrusions. On the other hand, we saw in Section 3.1 that coupling the ACR scheme to the STEN network could mitigate excitable behavior. In a chemotaxing cell, this would not be desirable, as the cell needs to move. However, if we can direct the effect of ACR in a spatially selective way, this could be beneficial. We now investigate whether the inclusion of an absolute concentration robustness module to the LEGI-STEN system could improve the chemotactic response. In particular, we propose to use the LEGI mechanism to direct the ACR control towards the rear of the cell.

As before, we use the back molecule as the target species for ACR, as this molecule is found preferentially at the back of the cell. Thus, as in Section 3.1, molecule B would be rendered robustly stable through ACR in both mean and variance. As B is directly involved in the mutually inhibitory loop of F, this would consequently allow us to control the firings of the system. A necessary component still missing, however, is a spatial regulator of the binding and unbinding reactions of ACR, i.e. something to make the parameters *α* and *β*, of Eqn. 7, spatially dependent. For this purpose, we use the local signal of the LEGI mechanism, E*. As this signal is found preferentially at the front of the cell, we use it to inhibit the two binding reactions of the ACR scheme:

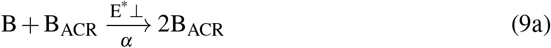

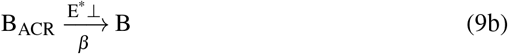

Here, E*⊥ refers to the inhibition of the reaction by the presence of *E*\*; specifically:

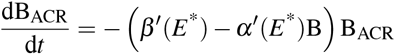

where

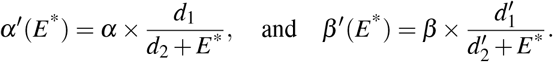

The overall scheme is illustrated in Fig. 7A. Fig. 7B shows a kymograph of a simulation of the combined LEGI-STEN-ACR network. For the first part of the simulation, the ACR mechanism is not active, owing to the low concentration of S. Upon addition of the chemoattractant (at *t* = 300), LEGI responds. Since the activation of E* is fast, the ACR is also switched on. Because E* is active near the front of the cell, we see how the action of ACR abrogates activity in the cell owing to the strict mean and variance control of B - similar to what we saw earlier. However, unlike the case of Fig. 6B, due to the spatial regulation of the ACR controller, i.e. owing to the inhibition of ACR at the front by E*, it is only at the rear that the firings disappear completely. Fig. 7C illustrates this in time by plotting line scans of the kymograph at positions “a” and “b”. While the front of the cell (point “a”) shows an increase in the number of firings (due to the LEGI mechanism), the back of the cell (point “b”) becomes completely quiescent because of the spatially-regulated ACR mechanism.

**Figure 7:**
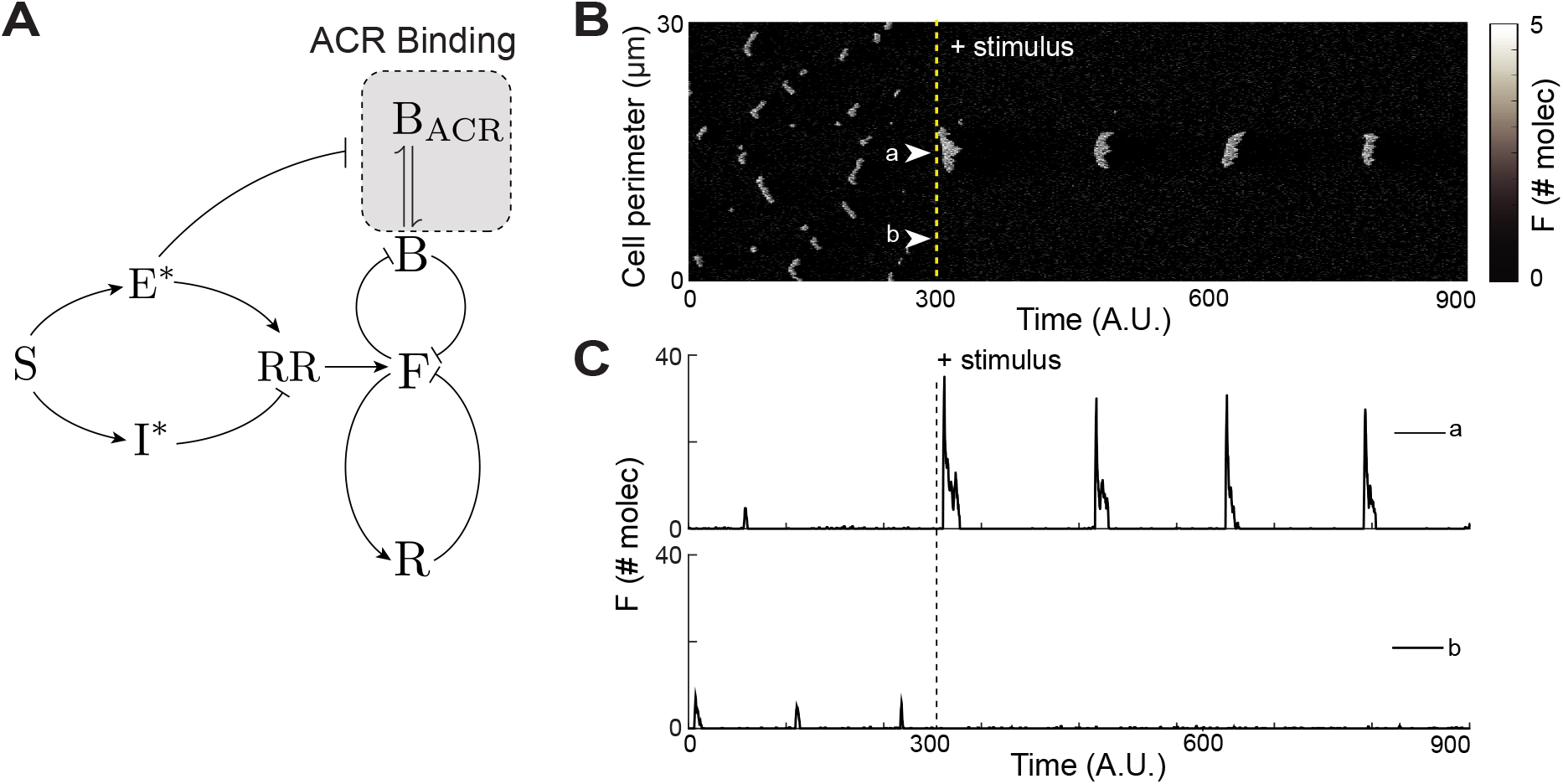
Implementation of ACR binding in LEGI-STEN system. (A) Schematic of the coupled LEGI-STEN-ACR system showing the incorporation of ACR binding (dashed box) through interaction between B and B_ACR_. (B,C) Kymograph and respective time profiles of F from line scans at locations a and b denoted by arrows. The gradient stimulus was applied at *t* = 300 A.U. at which point the ACR mechanism was also engaged.

The simulations above show a reduction in the activity of the excitable network. To test the effect of these firings on the chemotactic efficiency of the cell, we simulated the movement of the cells using a center of mass approximation. To this end, the activity of the variable R, as a function of the angle *θ* around the perimeter of the cell was translated into a vector normal to the cell surface. The vector sum was used as a measure of the net protrusive force (Fig. 8A). This net force was then appropriately scaled (so that it was in the range of experimentally observed protrusive pressures 0.5–5 nN/*μ*m^2^) and then fed into a viscoelastic model of *Dictyostelium* mechanics [53]. In this model, *F_x_*, the net force in the *x*-direction (the direction of the gradient) alters the center-of-mass position (CM_*x*_) through the following dynamics:

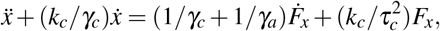

with a similar equation for the displacement in the *y*-direction (CM_*y*_). To illustrate the movement generated through this approximation, in Fig. 8B, we plot the trajectories of 10 cells migrating randomly in absence of any stimulus. We then simulated and compared the movement of 100 cells each with and without the ACR mechanism. While cells relying only on the LEGI-STEN coupling still moved along the direction of the chemoattractant source (Fig. 8C, top), the chemotactic efficiency was signficantly enhanced by the addition of ACR controller (Fig. 8C, bottom). A comparison of the final position in the *x*-direction shows that the ACR controller enabled the cells to travel further in the direction of needle (Fig. 8D. left), when compared to the cells without the ACR control. Moreover, the ACR mechanism also restricted their movements in the vertical direction as can be seen from the difference in the standard deviation of the final position along the *y*-direction, which was 1.45 *μ*m for the system without ACR and 0.68 *μ*m with (Fig. 8D, right).

**Figure 8:**
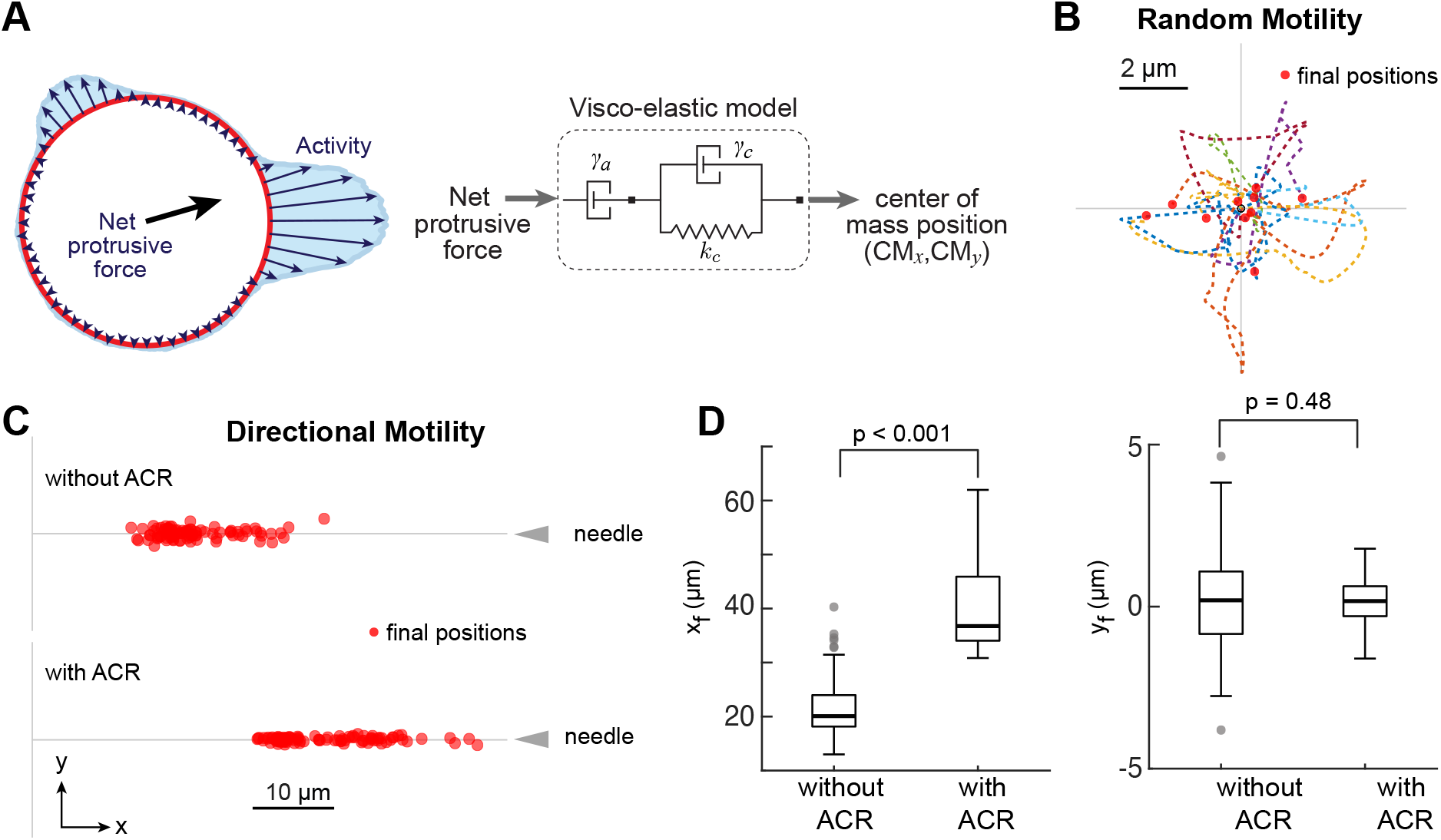
Comparison of cell motility. (A) The schematic shows the center-of-mass approximation scheme, where the net protrusive force was calculated from the R activity along the cell perimeter. This force was then fed into the viscoelastic model of a cell to obtain the displacement coordinates (CM_*x*_,CM_*y*_). (B) The trajectories of 10 cells undergoing random motility for a period of 100 time steps. The cells started at origin and the final positions are denoted with red circles. (C) Comparison of gradient sensing without (top) and with (bottom) ACR. The final positions of 100 cells are shown after 900 time steps. (D) Boxplots showing the distribution of the *x* and *y* coordinates of the final cell positions without and with ACR.

## 4 Discussion

For efficient chemotaxis, cells must overcome noisy chemical cues and intrinsic stochastic perturbations to interpret the chemotactic gradient and direct protrusions in the direction of the chemotactic cue. There are broadly two ways to enhance directional migration: increasing the protrusion frequency at the front of the cell or lowering the frequency at the back of the cell. Because protrusions are driven by the crossing of the threshold of the excitable system, lowering the threshold would have the effect of generating more protrusive activity. This has been demonstrated experimentally [10]. However, because of the refractory period of the excitable system, there is an underlying upper bound to the achievable frequency of firings. Moreover, lowering the threshold too far leads to a loss of stability which can cause oscillatory behavior [40] or permanently high levels of activity [41], neither of which results in motile cells.

For this reason, spatial control of the threshold to reduce the firings at the back of the cell is more promising: a perfectly quiescent back ensures that all the protrusive activity is directed towards the chemoattractant gradient. Cells achieve this spatial threshold regulation through the LEGI mechanism, which lowers the threshold at the front and raises it at the rear [28]. When combined with a synthetic increase of activity, spatial control of the threshold leads to fan-shaped cells that migrate 3-4 times faster than wild-type cells [40].

LEGI alone may not be enough to completely eliminate protrusions away from the source, particularly in shallow gradients. Even if the threshold is high at the back of the cell, fluctuations may still cause unwanted firings if these are sufficiently large. For this purpose, in this study, we considered how the efficiency of chemotaxis could be improved through the inclusion of a mechanism for regulating these fluctuations of various regulators of the chemotactic response. We saw that variance control of the stochastic inputs using absolute concentration robustness limited the number of firings of the excitable system (Fig. 4), by converting the species concentration distribution to a Poisson distribution. The variance of a Poisson process equals the mean, and, thus, this creates tight control not only over the species mean but also the variance.

This control, however, should only occur at the back of the cell. In this study, we not only show that ACR can control the number of firings of the cell, but also suggest a seamless method to incorporate ACR selectively at the rear without the need for an additional external biasing input. That is, by linking the ACR molecule with the excitatory species of LEGI we achieve back-quiescence simply on the addition of the chemoattractant gradient. Previously [61], we suggested that ACR be used to increase cell migration capability, but in that study the ACR controller was added externally without the use of the LEGI system. In contrast, by combining LEGI and ACR as we do here, we are proposing a feedforward control scheme where the chemoattractant gradient itself is sufficient to turn on ACR at the back of the cell and achieve better chemotaxis.

The controller we propose here has the advantage of being relatively simple as it relies on much of the existing architecture of the system. Incorporating this scheme experimentally only requires that we have a candidate for the excitation element of the cell and a protein that is inhibited by the local signal. The G*βγ* subunits, that couple receptor signaling to the downstream effectors, are the likely candidates for the LEGI exciter (E*) [15, 62, 63]. We next need a molecule B_ACR_ that can bind to the back markers, which include the phospholipids PI(4,5)P_2_ and PI(3,4)P_2_ [49], the phosphatase PTEN [64], the novel protein Callipygian [65] and the actin motor myosin II [66]. PI(4,5)P_2_ is inhibited by the phosphatases that degrade it (for example, Inp54p [40]), PLC [67] and the kinase PI3K [68]. Interestingly, the the activities of PLC and PI3K are upregulated by the receptor as would be required for the scheme suggested here. Thus, it is possible that cells may already be making use of a combined LEGI-STEN-ACR mechanism to direct their migration.

